# Males of a suboscine lekking bird exhibit neighbor-stranger discrimination

**DOI:** 10.64898/2026.10.15.759145

**Authors:** Josephine T. Tagestad, J. Patrick Kelley, Camilo Alfonso, Corey E. Tarwater

## Abstract

The ability to discriminate the vocalizations of neighbors from strangers is one way individuals can reduce conflict and minimize the associated physical and energetic costs. Although individual vocal variation and neighbor-stranger discrimination has been tested across various taxa, within birds, it is more commonly investigated in oscine passerines and territorial species, while suboscine passerines and lekking species are often neglected. Red-capped manakins, *Ceratopipra mentalis*, are a suboscine lekking bird that is known to exhibit individual variation in their whistle-peep vocalizations. We performed a playback experiment at *C. mentalis* leks to evaluate whether they exhibit neighbor-stranger discrimination at the lek-scale. We found that *C. mentalis* males responded more quickly to strangers compared to neighbors, suggesting that strangers are a greater threat to males than neighboring individuals. Neighbor-stranger discrimination was only observed in vocal responses, not movement responses, and differences in total vocal responses were only observed when we evaluated the changes in response to neighbors versus strangers over time. These findings add to a growing body of literature demonstrating that non-territorial and suboscine species are capable of neighbor-stranger discrimination. Further examination of neighbor-stranger discrimination in these understudied species can yield insights into the functions of vocalizations, and in particular male-male interactions and mate choice in highly sexually-selected species.

## INTRODUCTION

Across multiple taxa, the ability to discriminate between familiar “neighbors” and unfamiliar “strangers” allows individuals to avoid unnecessary conflicts and engage in aggression only when threatened, thereby optimizing expenditure of time and energy (Temeles 1990, Christensen and Radford 2018). A prerequisite to differentiating neighbors from strangers, called neighbor-stranger discrimination (hereafter NSD), is that the different communication signals that organisms use, such as calls, mechanical sounds, visual cues, or olfactory cues, must have a detectable signal of familiarity between individuals (Tripovich et al. 2008, Christensen and Radford 2018, Robertson 2018). If individuals can discriminate between conspecifics and if neighbors and strangers generate different responses from focal individuals, making the detection of NSD feasible, there are two alternative predictions (Bard et al. 2002). The first is a stronger response by focal individuals to the signal of stranger individuals, compared to familiar neighbors, dubbed the “Dear Enemy” effect (Fisher 1954). The second is a stronger response to neighbors compared to strangers, called the “Nasty Neighbor” effect (Müller and Manser 2007).

Dear Enemy effects typically occur when the relative threat of a stranger is greater than that of a neighbor, such as when territories and pairs are already established and stable (Briefer et al. 2008). In contrast, Nasty Neighbor effects tend to be more prevalent during settlement of territories or pair formation, when neighbors pose a significant threat (Briefer et al. 2008). Despite the importance of NSD for individual fitness, sexual selection, and conspecific interactions (Christensen and Radford 2018), it remains unclear whether most species are able to differentiate neighbors from strangers, and if they can, to which group they respond more strongly.

Studies of NSD in birds typically focus on oscine species. In contrast, suboscine birds are infrequently examined, despite representing approximately 20% of the world’s passerine species (Sibley and Monroe 1990, Oliveros et al. 2019). Suboscines generally do not exhibit song learning (see Kroodsma et al. 2013 for an exception) and have less complex syrinxes compared to oscines (Kroodsma 1984, Brenowitz 1991, Baptista and Trail 1992). These traits have led to the assumption that suboscines lack the physical complexity to exhibit individual variation in calls, and thus cannot discriminate between individuals based on vocal cues (Baptista and Trail 1992, Garcia et al. 2017). This has resulted in only a handful of NSD studies on suboscines in the last two decades, and these studies have yielded contrasting results (Westcott 1997, Bard et al. 2002, Lovell and Lein 2004, Wiley 2005).

Studies of NSD have also been generally limited to species with individuals or pairs defending a physical space from neighbor or stranger intruders (Christensen and Radford 2018). Nevertheless, the benefits of being able to discriminate among individuals is predicted to be equal or more important for group-living species (Getty 1987). Within this category, lekking species whose males gather in groups to display for females are particularly understudied, yet the ability to discriminate between individuals within one’s lek compared to individuals from outside the lek would be predicted to be strongly favored (Hale et al. 2014, Christensen and Radford 2018). In lekking systems, disruption of breeding activity is frequently disadvantageous to both male and female reproductive success, and discriminating between individuals to assess threats could therefore be beneficial to maximizing reproductive success (Foster 1983).

The red-capped manakin, *Ceratopipra mentalis*, is a suboscine passerine whose males are known to exhibit individual variation in their contact calls (Tagestad et al. in prep.). *C. mentalis* males congregate in leks and exhibit lek fidelity, returning to the same lek site yearly during the breeding season (Kricher and Davis 1998). In manakin species in general, there is often a dominance hierarchy among males, with minimal turnover in males among seasons and years (Ryder et al. 2011, Lukianchuk and Doucet 2014). Males benefit in long-term reproductive success by remaining in the same lek with a stable dominance hierarchy (Durães et al. 2008, DuVal 2013). Therefore, we predict males would benefit from the ability to discriminate between “neighboring” males in their own lek and “stranger” males from a different lek. We conducted a NSD experiment towards the end of the breeding season by broadcasting playbacks of neighbor and stranger calls at *C. mentalis* leks. We predicted that males would exhibit NSD and they would respond more aggressively to strangers than to neighbors, as late breeding season leks are more stable than early season leks (Théry 1990, Westcott 1997, Rosselli et al. 2002).

## METHODS

### Study species and sites

*C. mentalis* males exhibit a condensed exploded lek in which three to 10 males engage in uncoordinated display for females during the dry season and at least two months of the rainy season (Kricher and Davis 1998). Males are in visual or auditory contact and frequently perform a variety of calls, including whistle-peeps, short “peeps” unaccompanied by any whistle, long “whistle-screeches” that begin with a circuit flight off the perch and culminate in a screaming whistle and explosive screech, and mechanical noises produced by behaviors such as “wing-snaps” (described in Skutch 1949, Bostwick and Prum 2003).

We examined males in 13 leks at three sites across the Isthmus of Panama. Leks were a minimum of 140 m apart and each site was separated by at least seven kilometers and a significant spatial barrier (river, large multi-lane highway, and/or city). With this spatial separation and the lek fidelity observed in males, bird movement between leks and sites did not likely occur during the breeding season.

### Experiment setup

We previously found *C. mentalis* males exhibit individual variation in their whistle-peep calls (Tagestad et al. in prep.) and, accordingly, chose this vocalization as our playback stimulus. It was not possible to record data blind, as our study involved focal birds in the field. We spent five days (May 28-June 1, 2019) visiting the leks where we planned to perform a playback to record whistle-peeps from males at leks and to estimate the number of males attending leks. We recorded a minimum of four whistle-peeps at each lek, and performed an estimate of the number of males present, by listening for vocalizations to detect perch location and seeking visual confirmation for a minimum of 30 minutes. Male quantity estimates were confirmed as we returned to each lek for playback.

Initial whistle-peep calls were recorded using a Tascam™ DR-05 Linear PCM internal microphone (WAV 16 bit 44.1k stereo with input 80). We then extracted the three clearest whistle-peeps from each lek, selecting uninterrupted calls containing minimal ambient noise, and organized these into stimulus files with three evenly-spaced whistle-peeps. Lek activity is extremely variable, so we selected a moderate stimulus of three-per-minute that would realistically be encountered by males at this point in the breeding season. This was determined by assessing the average timing of whistle-peep calls recorded during our initial lek visit. We did not alter the order in which the whistle-peeps were initially recorded. Each stimulus file consisted of three whistle-peeps, each separated by 25 seconds of silence (see Fig. 1a). Playback files in .wav format were constructed using these stimulus files, with one minute of stimulus (first treatment) followed by 15 minutes of no broadcasting, followed by another minute (second treatment) of stimulus and another 15 minutes of silence (Fig. 1a). We randomized the order of stimulus type for each trial, resulting in eight trials presenting a stranger stimulus first and nine trials presenting a neighbor stimulus first. Each focal lek received both the neighbor and stranger treatments. We defined “neighbors” as birds present at the focal lek. We defined “strangers” as birds from a different site (3.55-23.35 km away). Every lek received whistle-peeps from a different male for the same treatment group (neighbor or stranger), but the same male whistle-peeps could be used for both a neighbor treatment and a stranger treatment on different leks.

**Figure 1.**
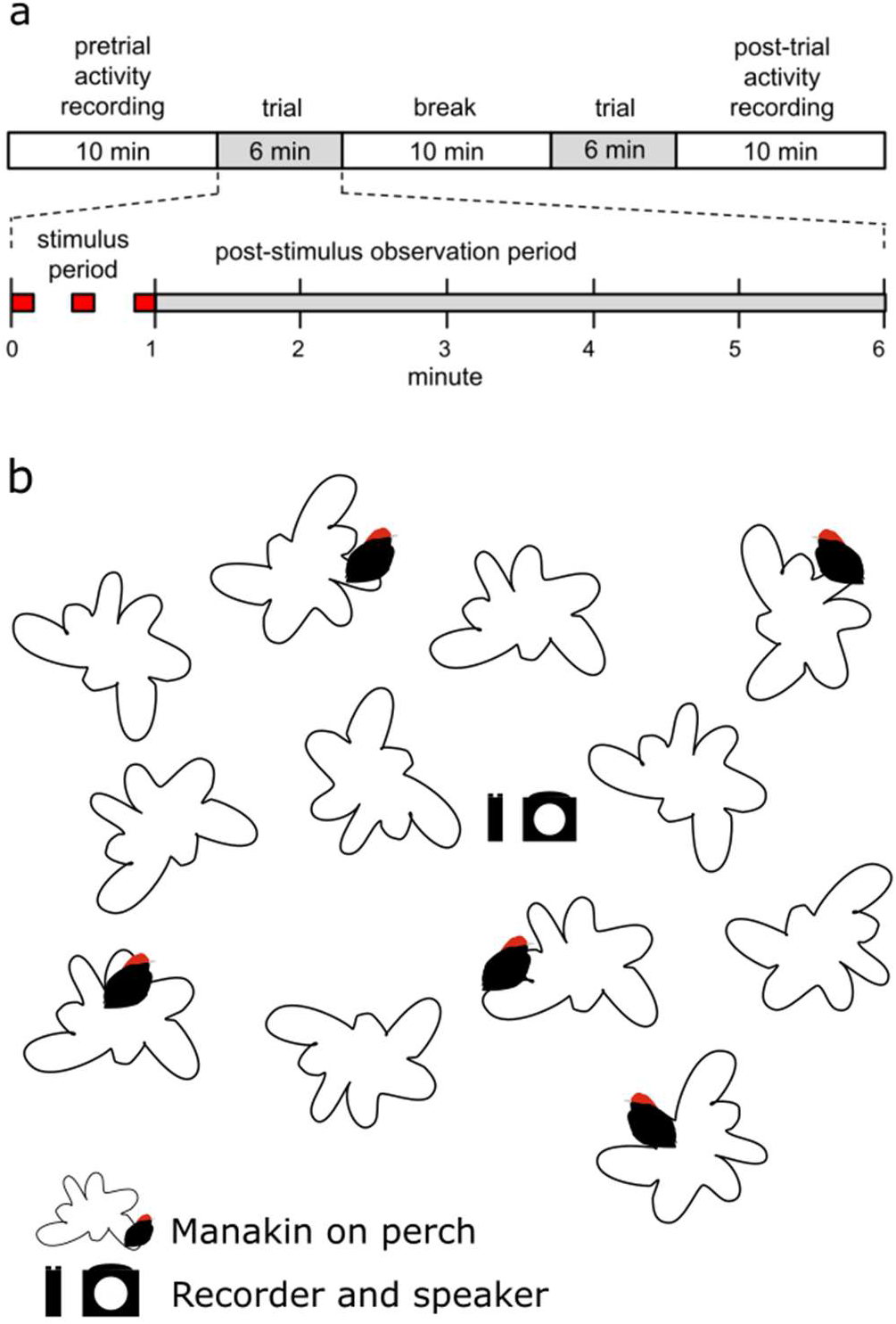
(a) Timing and order of each playback and trial. “Pretrial activity recording” and “Break” periods were silent (e.g, did not broadcast any noise). The “Stimulus” period broadcasted three evenly spaced whistle-peep calls from a single treatment, neighbor or stranger. The “Observation” period was also silent with no broadcasting and broken down into five periods of one minute each. Behaviors during the stimulus and observation periods, together called a trial, were used in the analysis. (b) Simple representation of the experiment setup. Playback was conducted from an approximately central location in each lek

### Playback execution

We performed playbacks towards the end of the breeding season, when the dominance hierarchy within leks tends to be well-established, as opposed to the early breeding season, when interactions may be in flux (Théry 1990, Widemo 1997, Rosselli et al. 2002). Between 2-19 June 2019, we conducted 17 playback trials at 13 leks. Playbacks at the same lek were conducted at least two days apart. At each lek, we passively recorded the general lek activity for a minimum of 10 minutes prior to the start of the trial to ensure that there were no large disruptions, such as predator arrival or sudden weather changes. We broadcast playback from an iPod Nano connected to a Pignose 7-100 portable amplifier placed in the approximate center of the lek (Fig. 1b). All audible responses (e.g. wing-snaps, peeps, and whistle-peeps) were recorded on a handheld Tascam™ DR-05 Linear PCM internal microphone (16-bit, 44.1k sampling rate, uncompressed WAV format). The timing and order of each playback and trial is shown in Fig. 1a. Trials were conducted and responses were passively recorded without vocal commentary by the observer beyond an initial metadata description. We evaluated the activity of the lek as a whole, rather than a single individual’s behavior, as focusing observations on a single bird would result in a skewed response depending on the individual’s position in the lek (e.g., dominant versus subordinate male).

### Behavioral responses

Responses were extracted from the playback recordings by a single person (JTT). For each recording, the one minute stimulus period and five minutes of post-stimulus observations were used to assess responses to treatments. Four different responses that encompass both the number of responses and the time it takes to respond to the stimulus were measured — total vocalizations, vocal latency, total movements, and movement latency; all responses are commonly used in playback studies to measure aggression (Bard et al. 2002, Lovell and Lein 2004, Wiley 2005, Seddon and Tobias 2007, Briefer et al. 2008, Skierczyński and Osiejuk 2009, Wilson and Mennill 2010, Koetz-Trowse et al. 2012, Draganoiu et al. 2013, Hale et al. 2014, Battiston et al. 2015, Cornec et al. 2015, Moskát et al. 2017).

Total vocalizations included the total number of songs and calls, with more vocalizations signifying a more intense response to the stimulus (Bard et al. 2002, Lovell and Lein 2004, Wiley 2005, Briefer et al. 2008, Skierczyński and Osiejuk 2009, Wilson and Mennill 2010, Hale et al. 2014, Battiston et al. 2015, Cornec et al. 2015). To quantify total vocalizations, we included the number of whistle-peep and peep calls. Whistle-peeps are the third most common vocalization from males at leks (based on data in Tagestad et al. in prep., 22.3% of 800 vocalizations) and serve as a contact call between adult males (Tagestad et al. in prep.). Peep calls are the most common vocalization given at lek (based on data in Tagestad et al. in prep., 40.0% of 800 vocalizations). The second most common vocalization (twitters) is more commonly given by a lek when a female or green-plumage juvenile male visits, whereas whistle-peep and peep calls are common calls exchanged between adult males. Vocal latency is the time it took for a bird to vocally respond to the start of the stimulus, and shorter latencies signify a more aggressive response (Lovell and Lein 2004, Seddon and Tobias 2007, Briefer et al. 2008, Koetz-Trowse et al. 2012, Battiston et al. 2015, Moskát et al. 2017).

Total movements include the number of flights and other visible movements, with increased movements by an individual signifying a more intense response to the stimulus (Bard et al. 2002, Lovell and Lein 2004, Wiley 2005, Seddon and Tobias 2007, Skierczyński and Osiejuk 2009, Draganoiu et al. 2013). To quantify total movements of all birds at the lek, we used two different sounds as a proxy for movement. First, any whistle-screech was recorded as a movement because 100% of all recorded whistle-screeches (based on data in Tagestad et al. in prep., n=30) were associated with a circuit flight. Second, any wing-snap was recorded as a movement. In 92.5% of 523 wing-snaps (based on data in Tagestad et al. in prep.), wing-snaps were associated with physical movement, such as a perch change, by the bird. Using these same data, we also confirmed that whistle-peeps and peeps were not strongly associated with movement (96% of 48 and 88% of 68 of these calls, respectively, were associated with no movement). For movement latency, we measured the time it took from the start of the stimulus to when any bird exhibited a movement response (Bard et al. 2002, Moskát et al. 2017). As with vocal latency, an earlier movement response signifies a more intense response to the stimulus (Briefer et al. 2008, Bradley et al. 2012).

### Analyses

All analyses were conducted in Program R 3.6.1 (R Core Team 2019) using generalized additive mixed models (GAMMs), to allow for nonlinear impacts of predictors, or generalized linear mixed models (GLMMs). We conducted separate analyses for each set of response variables: 1) total vocalizations and movements per minute of the trial and 2) vocal and movement latency responses over the course of the entire trial. Response variables were the following: total vocalizations (number of whistle-peeps and peeps), vocal latency (number of seconds from the start of the stimulus until a peep or whistle-peep), total movements (probability of movement, rather than number of movements owing to many 0s in the data), and movement latency (number of seconds from the start of the stimulus until a movement). We compared, using AICc, the random effects of lek, plot, and site (as three separate random effects owing to too small sample sizes to allow for hierarchical random effects), and found that using lek as the random effect was the best model. To control for potential variation in responses due to nuisance variables, we included time of day the experiment started, day in June of the experiment, number of males in a lek, and order of stimuli presented to a lek (either neighbor or stranger as the first treatment) as fixed effects in the models. Model results were based on the global model for each response variable (no model selection was conducted). R^2^ were calculated using the conditional R^2^.

#### Vocalizations and movements per minute (n=204)

Total vocalizations and movements were analyzed per minute of the trial (six minutes in total; one minute of stimulus and five minutes of post-playback) to examine if responses changed between neighbor and stranger treatments over the course of the playback. GAMMs were used for both analyses: a Poisson error distribution (log link) was used for the vocal response, and a binomial error distribution (logit link) was used for the movement response. We first tested which variables required smoothers (were nonlinear) and retained smoothers only for nonlinear variables (minute and day for vocal response and minute, day, and hour of the day for the movement response). Estimated degrees of freedom (e.d.f.) in GAMMs represent how nonlinear a covariate is; an e.d.f. of 1 represents a linear relationship and, as e.d.f. increases, nonlinearity increases. Fixed effects include treatment (neighbor or stranger), order of stimuli presented to a lek, time of day the experiment started, day in June of the experiment, number of males in a lek, and the interactions between minute of the experiment and treatment.

#### Latency to respond (n=34)

We used GLMMs with a negative binomial error structure to model latency to vocalize and move over the entire experiment. As above, we first tested whether there were any nonlinear effects, but only tested for quadratic effects owing to the small sample size. Further, due to this small sample size, we could not have more than four fixed effects in the model at one time. Therefore, we tested whether our nuisance variables influenced latency, and if they did not, they were removed from the global model. To model latency to vocalize, we retained treatment, hour of the day, order of stimuli, and number of males in the global model, and day of the experiment was removed. To model latency to move, we retained treatment, hour of the day, number of males, and a quadratic effect of number of males in the global model, and day of the experiment and order were removed.

## RESULTS

*C. mentalis* males responded differently to neighbors versus strangers vocally, both in terms of total vocalizations and latency, but they did not respond differently in terms of movement (Tables 1 and 2). For total vocalizations in response to a stranger, males produced the highest number of vocalizations when the stimulus was still going (first minute of the trial), and then response slowly declined over time (Fig. 2). In response to a neighbor stimulus, males responded the highest in the middle of the trial, and produced the fewest vocalizations when the stimulus was still going (Fig. 2). If we had examined only the number of vocalizations in response to neighbors and strangers across the entire trial, rather than per-minute, we would have found no differences in response between them (see Table 1a linear term of treatment). It was only when we examined responses over time that we observed a different response (see Table 1a nonlinear term of the interaction between minute and treatment). Hour of the day and day of the playback also impacted the response of individuals to the treatment (Table 1a). Males produced more vocalizations later in the day and response to stimuli was lower earlier and later in June compared to the middle of June (Table 1a). Given that p-values in GAMMs are approximate and that the nonlinear relationships we found were quadratic, we re-ran the analysis with a GLMM (using the same parameters) and including quadratic relationships for the non-linear trends, and obtained a p-value of 0.02 for the interaction between minute and treatment. Thus results are robust, regardless of type of model.

**Table 1.** Model results based on GAMMs for the total vocalization and movement models analyzed per-minute. Traits in bold indicate significant p-values (p<0.05)

| Model | Linear or nonlinear terms | Trait | Beta estimate or edf | SE | p-value |
| --- | --- | --- | --- | --- | --- |
| A) Number of vocalizations ($R^2 = 0.57$) | | | | | |
|  | Linear | intercept - treatment neighbor | -2.784 | 0.932 | 0.003 |
|  |  | order of treatment | -0.526 | 0.346 | 0.131 |
|  |  | **hour of day** | **0.230** | **0.065** | **0.001** |
|  |  | treatment- stranger | -0.099 | 0.098 | 0.314 |
|  |  | number of males | 0.213 | 0.113 | 0.062 |
|  | Nonlinear | minute X treatment - neighbor | 2.007 | 2.007 | 0.131 |
|  |  | **minute X treatment - stranger** | **1.000** | **1.000** | **0.013** |
|  |  | **day of playback** | **2.637** | **2.637** | **0.021** |
| B) Probability of movement ($R^2 = 0.30$) | | | | | |
|  | Linear | intercept - treatment neighbor | -2.796 | 1.549 | 0.073 |
|  |  | order of treatment | -0.607 | 0.470 | 0.199 |
|  |  | treatment - stranger | -0.294 | 0.324 | 0.364 |
|  |  | **number of males** | **1.219** | **0.501** | **0.016** |
|  |  | **number of males<sup>2</sup>** | **0.106** | **0.043** | **0.014** |
|  | Nonlinear | minute X treatment - neighbor | 1.000 | 1.000 | 0.366 |
|  |  | minute X treatment - stranger | 1.000 | 1.000 | 0.603 |
|  |  | **hour of day** | **1.618** | **1.618** | **0.004** |
|  |  | **day of playback** | **2.138** | **2.138** | **0.006** |

**Table 2.** Model results based on GLMMs from vocal and movement latency models analyzed across the entire trial. Traits in bold indicate significant p-values (p<0.05)

| Model | Trait | Beta estimate | SE | p-value |
| --- | --- | --- | --- | --- |
| A) Latency to vocalize ($R^2 = 0.24$) | | | | |
|  | intercept -treatment neighbor | 5.608 | 1.270 | <0.001 |
|  | **treatment - stranger** | **-0.719** | **0.334** | **0.032** |
|  | **hour of day** | **-0.263** | **0.126** | **0.037** |
|  | order of treatment | -0.032 | 0.317 | 0.921 |
|  | number of males | 0.088 | 0.078 | 0.259 |
| B) Latency to move ($R^2 = 0.21$) | | | | |
|  | intercept -treatment neighbor | 6.725 | 1.501 | <0.001 |
|  | treatment - stranger | 0.537 | 0.363 | 0.14 |
|  | hour of day | -0.086 | 0.149 | 0.563 |
|  | **number of males** | **-0.692** | **0.318** | **0.029** |
|  | **number of males<sup>2</sup>** | **0.061** | **0.025** | **0.017** |

**Fig. 2.**
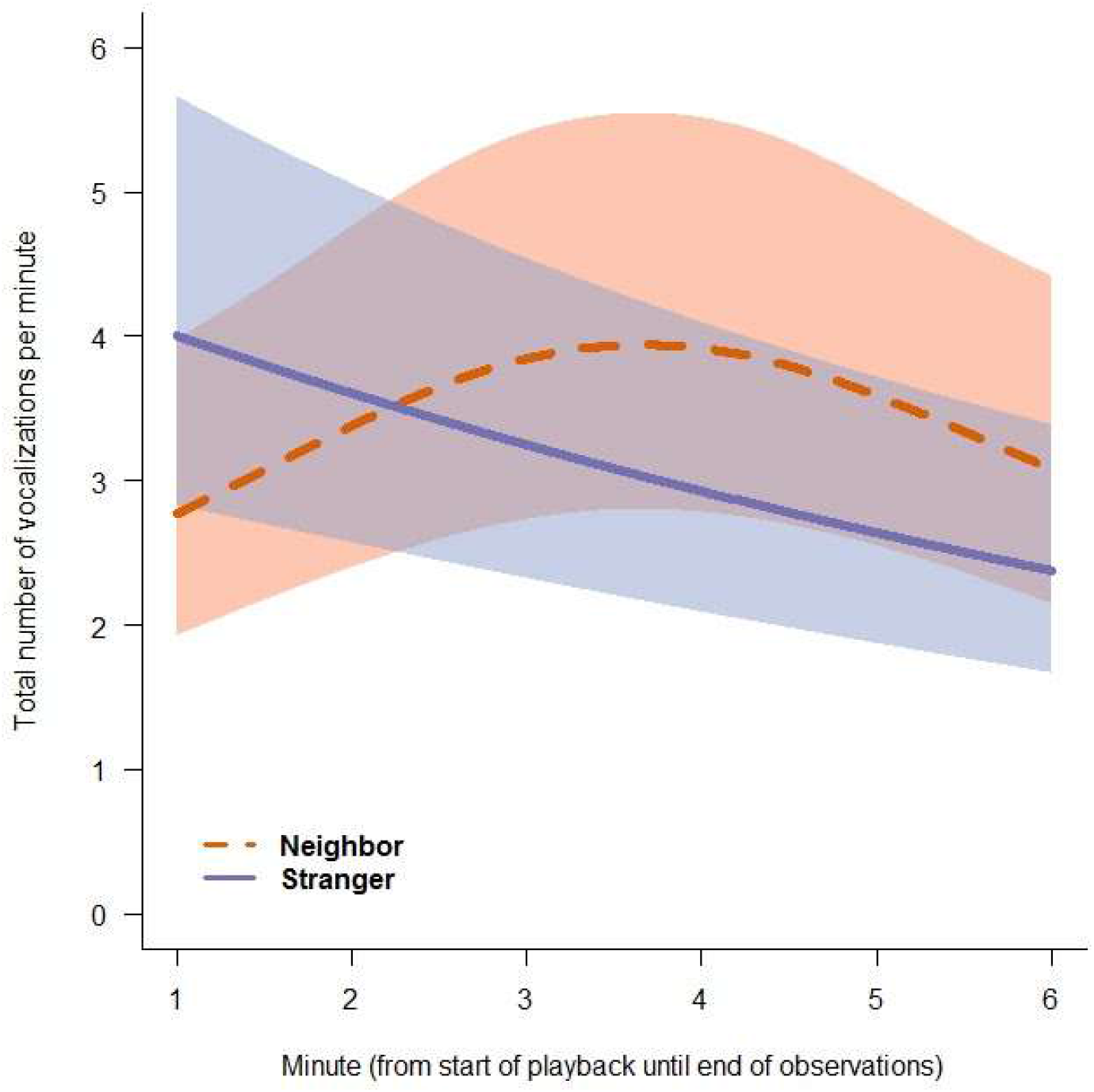
Total number of vocalizations produced by males in response to neighbor and stranger stimuli during each minute of the trial. Model predicted output based on GAMMs are graphed and SEs are shown. For the x axis, each minute marker corresponds to the total vocalizations in the preceding minute, such that minute 1 refers to the total vocalizations between minutes 0 and 1, minute 2 refers to the total vocalizations between minutes 1 and 2, minute 3 refers to the total vocalizations between minutes 2 and 3, etc

When evaluating latency to vocally respond, males responded more than twice as fast to the stranger stimulus than to the neighbor stimulus, consistent with our findings above indicating that males responded with more vocalizations to the stranger stimulus earlier in the trial (Table 2a, Fig. 3). Hour of the day also influenced response, with birds responding more quickly earlier in the day (Table 2a).

**Fig. 3.**
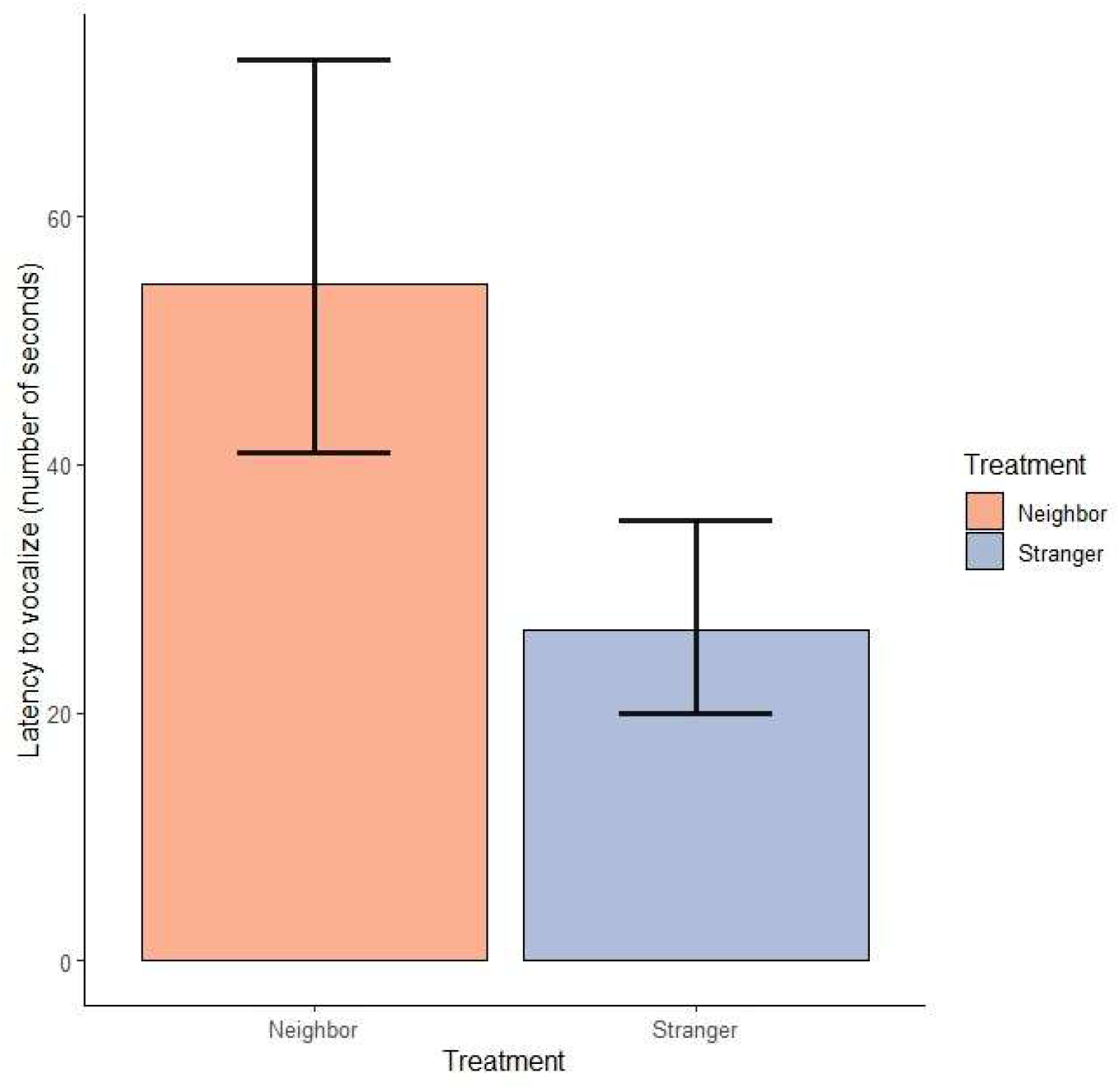
The latency of males to vocalize (number of seconds between the start of the stimuli and first vocal response) in response to neighbor and stranger stimuli. Model predicted output based on GLMMs are graphed and SEs are shown

For movement responses, treatment did not influence the probability of movement per-minute (Table 1b) or how quickly the males moved after the start of the stimulus (Table 2b). Males were more likely to move in response to playbacks later in the day and in the middle of June, compared to earlier and later in June (Table 1b). Thus, changes in daily activity and the time in the breeding season influenced how much males moved. Furthermore, the number of males at the lek influenced both the probability of movement and latency to move, with individuals being more likely to move and move faster with an average number of males, compared to few or many males in the lek (Tables 1b and 2b).

## DISCUSSION

By experimentally testing NSD in a bird species, we found four key results. First, we showed males exhibit NSD in a lekking suboscine. This finding adds to a growing body of literature demonstrating that suboscines are capable of NSD, despite being less vocally complex than oscines, and that NSD does occur in non-territorial species. Second, males had a stronger response to strangers compared to neighbors, consistent with the “Dear Enemy” hypothesis. This suggests that strangers are a greater threat to males than individuals within the same lek. Third, NSD was only observed in male vocal responses (latency to respond and total vocalizations), not in their movements. This may arise due to greater energetic costs and risks associated with movement, compared to vocalizations, or methodological constraints. Fourth, differences in total vocalizations were only observed when we evaluated the changes in response to neighbors versus strangers over time, highlighting the need for studies to use similar approaches to understand responses. Overall, suboscine NSD is generally understudied, particularly in lekking birds, and investigating NSD can yield a better understanding of sexual selection and social behaviors in these species.

### NSD in lekking suboscines

Our study and a handful of others on suboscines suggest that vocal complexity and song learning, as found in oscines, are not prerequisites to being able to distinguish familiar and unfamiliar conspecifics, despite this long-held assumption (Kroodsma 1984, Kroodsma and Konishi 1991, Baptista and Trail 1992, Stoddard 1996, Lovell and Lein 2004, Koloff and Mennill 2011, Garcia et al. 2017). Instead, vocal discrimination remains possible in any species that exhibits vocal characteristics that vary consistently among individuals (Falls 1982). *C. mentalis* is a suboscine that exhibits individual variation in vocalizations (Tagestad et al. in prep.) and can discriminate between familiar and unfamiliar conspecifics. We suggest further studies of suboscines will continue to break down long-held assumptions, and we argue there remains a great need for studies on the ecological, behavioral, and evolutionary impacts of NSD in subsocines.

Studies of NSD in lekking species are extremely limited, and here we present the first evidence of NSD in a lekking manakin species. The two previous studies of NSD in lekking species have found contrasting results, with no NSD present in the boom calls of greater prairie chickens, a lekking species with delineated territories, and NSD observed in ochre-bellied flycatchers, a species that defends both solitary territories and delineated lek territories (Westcott 1997, Hale et al. 2014). Here we evaluated the response of the lek as a whole, comparable to studies of group-living species, rather than looking at an individual’s capability to discriminate between neighbors and strangers, as done in previous lekking species (Westcott 1997, Hale et al. 2014, Christensen and Radford 2018). NSD in group-living species is often found to be highly consequential, as territorial conflict in these species can include not just effects on the individual, such as risk of injury and loss of time, but also effects on the group, such as group dynamics and behavioral interactions leading to reproductive success (Christensen and Radford 2018).

NSD may be important in establishing and maintaining the dominance hierarchy within a lek, an essential aspect of manakin social dynamics that strongly influences mating success (Brodt et al. 2014, Lukianchuk and Doucet 2014, DuVal et al. 2018). Dominance hierarchies minimize aggressive interactions, and NSD can potentially aid with the establishment and maintenance of hierarchies among males (Beehler and Foster 1988). Thus, we expect that in lekking species, just like in other territorial and group-living species, minimization of aggressive interactions to reduce loss of energy and risk of injury drives NSD (Temeles 1994, Christensen and Radford 2018). We suggest that recognizing individuals within the same lek would be beneficial in optimizing energy expenditure, allowing males to minimize aggressive interactions when, for example, competing for females. Male discriminatory abilities are likely to have an impact on social interactions, and in strongly sexually-selected species with high mating skew, the ability of both males and females to discriminate among individuals may play a key role in sexual selection. Furthermore, individually distinct vocalizations frequently carry quality information, and females may be able to recognize and utilize vocalizations to discriminate between potential mates (Bard et al. 2002, Cornec et al. 2015, DuVal et al. 2018). Thus, further investigation of the role that NSD plays in sexual selection and social interactions in lekking species is sorely needed.

### Dear enemy effects

*C. mentalis* male response was consistent with the Dear Enemy effect, whereby males responded more quickly to the calls of strangers compared to neighbors. The presence of Dear Enemy effects suggests that conspecifics that are not part of the lek (e.g., strangers) pose a more significant threat to the established lek compared to a different lek member (Jaeger 1981). Males outside of established leks tend to be juveniles searching for mating opportunities and leks to join, as *C. mentalis* is not known to have floater (e.g., no lek) adult males. Since adult males are known to exhibit lek fidelity (Kricher and Davis 1998), and the same males will interact frequently across breeding seasons, newcomers are clearly threatening to the stability of the existing lek dynamics. In some cases, the Dear Enemy effect is observed to be present during times of stability in territorial boundaries, but not when territories are in flux (Briefer et al. 2008, Christensen and Radford 2018). Given that manakin males often return to the same lek year after year and dominance hierarchies can persist across years, we may be less likely to see long-term temporal variation in their response to neighbors versus strangers; however, further research is needed.

### NSD in vocalizations versus movement

*C. mentalis* males exhibited NSD in vocalizations, whereas we did not detect a different response in the probability of moving or the latency to move. There are a number of possible reasons for these findings. First, movement is energetically more costly and riskier in terms of injury compared to vocalizations, and vocalizations alone often remain a reliable way to signal aggressive intent (Enquist 1985, Goldstein 1988). Thus, movement may be more likely to occur when an intruder is visible and there is a direct, nearby threat, whereas vocalizations may be more common in response to unseen intruders (Chamberlain and Cornwell 1971, Smith and Smith 2000). Furthermore, manakin males, like other lekking males, already expend huge amounts of energy displaying (Widemo 1997, Barske et al. 2014), likely increasing the benefits of responding only vocally. Second, males may have responded in terms of movement, but they did so similarly between neighbors and strangers. The finding of no differences in movement does not mean that males did not move, only that they moved similarly across treatments. Lastly, the sounds used as a movement proxy may not adequately represent how much individuals moved (e.g., they could have been moving when not vocalizing) or how they respond to an intruder. For example, the closest approach to a speaker is a common variable in NSD studies and may be a better indicator of aggression than the probability of movement alone (Bard et al. 2002, Lovell and Lein 2004, Seddon and Tobias 2007, Skierczyński and Osiejuk 2009, Draganoiu et al. 2013, Hale et al. 2014, Battiston et al. 2015). Few studies of bird species have found no differences in movement responses (see Koetz-Trowse et al. 2012 for an exception), and whether the lack of response we found was due to behaviors associated with displaying, such as higher energetic costs, or was methodological, requires further study.

### Changes over time in response to neighbors versus strangers

We found the total number of vocalizations differed between neighbors and strangers in *C. mentalis* only by examining changes in response over time. If we had only examined the number of vocalizations over the entire trial, we would not have found differences. These results suggest that analyzing behaviors over time yields a more comprehensive understanding of how individuals respond to threats. For example, by studying changes over time, we were able to determine the greatest response to strangers was at the beginning of the trial, while the greatest response to neighbors was in the middle of the trial. Males vocalized more in response to strangers compared to neighbors in the beginning of the trial, further supported by our findings that the latency to respond vocally was shorter with strangers. Thus, the stranger stimulus elicited a quick, intense response, but the response was not sustained. Therefore, in species where NSD responses are short-lived, analyses that do not examine responses at shorter temporal scales, as commonly occurs, may not detect NSD (for example, Bard et al. 2002, Wiley 2005, Hale et al. 2014, Battiston et al. 2015). Further examinations of NSD should analyze both changes over time and overall responses in an effort to create the most comprehensive understanding of response differences.

In conclusion, we found NSD in a lekking suboscine species, with males vocalizing more quickly in response to strangers compared to neighbors. Differences in vocalizations were only observed on a per-minute basis, suggesting that an examination of response over time is necessary to fully assess the NSD capabilities of any species. We argue that more studies are needed in lekking and suboscine species to fully address the ecological, evolutionarily, and behavioral implications of NSD.

## ACKNOWLEDGEMENTS

We thank the Smithsonian Tropical Research Institute (STRI) and the Wyoming Research Scholars Program for their financial and logistical support. All research was conducted under the auspices of permits provided by the Ministerio del Ambiente (SE/A-41-19) and approved under the Institutional Animal Care and Use committees at STRI (2019-0105-2022) and the University of Wyoming (# 20190522CT00368-01).

