## Supplemental Table 1 for "Males of a suboscine lekking bird exhibit neighbor-stranger discrimination"

The following tables include our data from the NSD trials for this project. It is broken into two separate tables: Trial Overall, which includes all variables from each trial as a whole, and Trial By Minute, which includes all variables from each minute of each trial.

Data: Trial Overall

| day in June | hour in day | site | plot_withinsite | lek | playback order | number of males | treatment | latency to vocalize | latency to move |
|---|---|---|---|---|---|---|---|---|---|
| 11 | 7 | 1 | 13 | 1 | NS | 7 | N | 33 | 321 |
| 11 | 7 | 1 | 13 | 1 | NS | 7 | S | 63 | 3 |
| 12 | 8 | 2 | 21 | 2 | SN | 1 | S | 18 | 57 |
| 12 | 8 | 2 | 21 | 2 | SN | 1 | N | 24 | 360 |
| 12 | 7 | 2 | 21 | 3 | NS | 8 | N | 0 | 39 |
| 12 | 7 | 2 | 21 | 3 | NS | 8 | S | 12 | 13 |
| 14 | 9 | 2 | 21 | 4 | NS | 10 | N | 100 | 112 |
| 14 | 9 | 2 | 21 | 4 | NS | 10 | S | 13 | 360 |
| 16 | 7 | 2 | 21 | 4 | SN | 10 | S | 24 | 360 |
| 16 | 7 | 2 | 21 | 4 | SN | 10 | N | 360 | 360 |
| 10 | 8 | 2 | 22 | 5 | SN | 5 | S | 42 | 173 |
| 10 | 8 | 2 | 22 | 5 | SN | 5 | N | 55 | 53 |
| 10 | 7 | 2 | 22 | 6 | NS | 5 | N | 44 | 5 |
| 10 | 7 | 2 | 22 | 6 | NS | 5 | S | 66 | 8 |
| 13 | 10 | 2 | 23 | 7 | SN | 8 | S | 6 | 44 |
| 13 | 10 | 2 | 23 | 7 | SN | 8 | N | 23 | 58 |
| 5 | 10 | 3 | 31 | 8 | NS | 7 | N | 143 | 81 |
| 5 | 10 | 3 | 31 | 8 | NS | 7 | S | 7 | 65 |
| 4 | 9 | 3 | 32 | 9 | NS | 5 | N | 8 | 45 |
| 4 | 9 | 3 | 32 | 9 | NS | 5 | S | 39 | 305 |
| 6 | 7 | 3 | 32 | 10 | SN | 7 | S | 25 | 76 |
| 6 | 8 | 3 | 32 | 10 | SN | 7 | N | 44 | 43 |
| 18 | 10 | 3 | 32 | 10 | SN | 7 | S | 33 | 360 |
| 18 | 10 | 3 | 32 | 10 | SN | 7 | N | 5 | 10 |
| 5 | 8 | 3 | 32 | 11 | NS | 6 | N | 68 | 73 |
| 5 | 8 | 3 | 32 | 11 | NS | 6 | S | 47 | 52 |
| 4 | 7 | 3 | 33 | 12 | SN | 8 | S | 46 | 360 |
| 4 | 7 | 3 | 33 | 12 | SN | 8 | N | 20 | 11 |
| 18 | 11 | 3 | 33 | 12 | SN | 8 | S | 7 | 47 |
| 18 | 11 | 3 | 33 | 12 | SN | 8 | N | 32 | 31 |
| 6 | 9 | 3 | 33 | 13 | NS | 7 | N | 6 | 14 |
| 6 | 9 | 3 | 33 | 13 | NS | 7 | S | 4 | 24 |
| 18 | 8 | 3 | 33 | 13 | NS | 7 | N | 39 | 46 |
| 18 | 8 | 3 | 33 | 13 | NS | 7 | S | 16 | 18 |

Data: Trial By Minute

| lek | playback order | number of males | playback id | treatment | minute of playback | latency to vocalize | latency to move | total movements | move (0/1) | total number vocalizations |
|---|---|---|---|---|---|---|---|---|---|---|
| 1 | NS | 7 | 06.12.131 | N | 3 | 33 | 321 | 0 | 0 | 0 |
| 4 | SN | 10 | 06.17.213 | S | 2 | 24 | NA | 0 | 0 | 0 |
| 4 | SN | 10 | 06.17.213 | S | 4 | 24 | NA | 0 | 0 | 0 |
| 4 | SN | 10 | 06.17.213 | S | 5 | 24 | NA | 0 | 0 | 0 |
| 12 | SN | 8 | 06.05.331 | S | 5 | 46 | NA | 0 | 0 | 0 |
| 4 | SN | 10 | 06.17.213 | N | 1 | NA | NA | 0 | 0 | 0 |
| 4 | SN | 10 | 06.17.213 | N | 2 | NA | NA | 0 | 0 | 0 |
| 4 | SN | 10 | 06.17.213 | N | 3 | NA | NA | 0 | 0 | 0 |
| 4 | SN | 10 | 06.17.213 | N | 4 | NA | NA | 0 | 0 | 0 |
| 4 | SN | 10 | 06.17.213 | N | 5 | NA | NA | 0 | 0 | 0 |
| 4 | SN | 10 | 06.17.213 | N | 6 | NA | NA | 0 | 0 | 0 |
| 12 | SN | 8 | 06.05.331 | N | 6 | 19.5 | 11 | 0 | 0 | 0 |
| 13 | NS | 7 | 06.19.332 | N | 5 | 39 | 46 | 0 | 0 | 0 |
| 13 | NS | 7 | 06.19.332 | N | 2 | 39 | 46 | 0 | 0 | 0 |
| 2 | SN | 1 | 06.13.210 | S | 2 | 18 | 57 | 0 | 0 | 0 |
| 2 | SN | 1 | 06.13.210 | S | 4 | 18 | 57 | 0 | 0 | 0 |
| 2 | SN | 1 | 06.13.210 | S | 5 | 18 | 57 | 0 | 0 | 0 |
| 2 | SN | 1 | 06.13.210 | S | 6 | 18 | 57 | 0 | 0 | 0 |
| 11 | NS | 6 | 06.06.324 | N | 1 | 68 | 73 | 0 | 0 | 0 |
| 11 | NS | 6 | 06.06.324 | N | 3 | 68 | 73 | 0 | 0 | 0 |
| 11 | NS | 6 | 06.06.324 | N | 4 | 68 | 73 | 0 | 0 | 0 |
| 11 | NS | 6 | 06.06.324 | N | 6 | 68 | 73 | 0 | 0 | 0 |
| 5 | SN | 5 | 06.11.2213 | N | 1 | 54.5 | 52.5 | 0 | 0 | 0 |
| 5 | SN | 5 | 06.11.2213 | N | 3 | 54.5 | 52.5 | 0 | 0 | 0 |
| 5 | SN | 5 | 06.11.2213 | N | 4 | 54.5 | 52.5 | 0 | 0 | 0 |
| 5 | SN | 5 | 06.11.2213 | N | 5 | 54.5 | 52.5 | 0 | 0 | 0 |
| 5 | SN | 5 | 06.11.2213 | N | 6 | 54.5 | 52.5 | 0 | 0 | 0 |
| 11 | NS | 6 | 06.06.324 | S | 5 | 46.5 | 52 | 0 | 0 | 0 |
| 11 | NS | 6 | 06.06.324 | S | 2 | 46.5 | 52 | 0 | 0 | 0 |
| 2 | SN | 1 | 06.13.210 | N | 5 | 23.5 | NA | 0 | 0 | 0 |
| 2 | SN | 1 | 06.13.210 | N | 6 | 23.5 | NA | 0 | 0 | 0 |
| 4 | NS | 10 | 06.15.213 | N | 1 | 100 | 112 | 0 | 0 | 0 |
| 4 | NS | 10 | 06.15.213 | S | 3 | 12.5 | NA | 0 | 0 | 0 |
| 4 | NS | 10 | 06.15.213 | S | 4 | 12.5 | NA | 0 | 0 | 0 |
| 4 | NS | 10 | 06.15.213 | S | 5 | 12.5 | NA | 0 | 0 | 0 |
| 4 | NS | 10 | 06.15.213 | S | 2 | 12.5 | NA | 0 | 0 | 0 |
| 9 | NS | 5 | 06.05.321 | N | 6 | 7.5 | 44.5 | 0 | 0 | 0 |
| 9 | NS | 5 | 06.05.321 | S | 3 | 39 | 305 | 0 | 0 | 0 |
| 9 | NS | 5 | 06.05.321 | S | 4 | 39 | 305 | 0 | 0 | 0 |
| 9 | NS | 5 | 06.05.321 | S | 5 | 39 | 305 | 0 | 0 | 0 |
| 8 | NS | 7 | 06.06.311 | N | 1 | 142.5 | 81 | 0 | 0 | 0 |

| | | | | | | | | | | |
|---|---|---|---|---|---|---|---|---|---|---|
| 10 | SN | 7 | 06.19.322 | S | 3 | 32.5 | NA | 0 | 0 | 0 |
| 10 | SN | 7 | 06.19.322 | S | 5 | 32.5 | NA | 0 | 0 | 0 |
| 10 | SN | 7 | 06.19.322 | S | 6 | 32.5 | NA | 0 | 0 | 0 |
| 10 | SN | 7 | 06.19.322 | S | 2 | 32.5 | NA | 0 | 0 | 0 |
| 8 | NS | 7 | 06.06.311 | S | 1 | 7 | 65 | 0 | 0 | 0 |
| 8 | NS | 7 | 06.06.311 | S | 6 | 7 | 65 | 0 | 0 | 0 |
| 10 | SN | 7 | 06.19.322 | N | 2 | 5 | 9.5 | 0 | 0 | 0 |
| 6 | NS | 5 | 06.11.222 | N | 6 | 44 | 4.5 | 1 | 1 | 0 |
| 1 | NS | 7 | 06.12.131 | S | 1 | 63 | 3 | 1 | 1 | 0 |
| 2 | SN | 1 | 06.13.210 | S | 3 | 18 | 57 | 1 | 1 | 0 |
| 11 | NS | 6 | 06.06.324 | N | 2 | 68 | 73 | 1 | 1 | 0 |
| 9 | NS | 5 | 06.05.321 | S | 6 | 39 | 305 | 1 | 1 | 0 |
| 8 | NS | 7 | 06.06.311 | S | 2 | 7 | 65 | 1 | 1 | 0 |
| 8 | NS | 7 | 06.06.311 | S | 3 | 7 | 65 | 1 | 1 | 0 |
| 8 | NS | 7 | 06.06.311 | S | 4 | 7 | 65 | 1 | 1 | 0 |
| 6 | NS | 5 | 06.11.222 | N | 5 | 44 | 4.5 | 2 | 1 | 0 |
| 6 | NS | 5 | 06.11.222 | S | 6 | 65.5 | 8 | 2 | 1 | 0 |
| 5 | SN | 5 | 06.11.2213 | S | 4 | 42 | 173 | 2 | 1 | 0 |
| 8 | NS | 7 | 06.06.311 | N | 6 | 142.5 | 81 | 2 | 1 | 0 |
| 8 | NS | 7 | 06.06.311 | N | 2 | 142.5 | 81 | 2 | 1 | 0 |
| 1 | NS | 7 | 06.12.131 | S | 6 | 63 | 3 | 4 | 1 | 0 |
| 10 | SN | 7 | 06.19.322 | N | 3 | 5 | 9.5 | 6 | 1 | 0 |
| 6 | NS | 5 | 06.11.222 | S | 1 | 65.5 | 8 | 7 | 1 | 0 |
| 6 | NS | 5 | 06.11.222 | N | 3 | 44 | 4.5 | 10 | 1 | 0 |
| 6 | NS | 5 | 06.11.222 | N | 2 | 44 | 4.5 | 11 | 1 | 0 |
| 1 | NS | 7 | 06.12.131 | N | 4 | 33 | 321 | 0 | 0 | 1 |
| 1 | NS | 7 | 06.12.131 | N | 5 | 33 | 321 | 0 | 0 | 1 |
| 4 | SN | 10 | 06.17.213 | S | 1 | 24 | NA | 0 | 0 | 1 |
| 4 | SN | 10 | 06.17.213 | S | 3 | 24 | NA | 0 | 0 | 1 |
| 4 | SN | 10 | 06.17.213 | S | 6 | 24 | NA | 0 | 0 | 1 |
| 12 | SN | 8 | 06.05.331 | S | 1 | 46 | NA | 0 | 0 | 1 |
| 12 | SN | 8 | 06.05.331 | S | 2 | 46 | NA | 0 | 0 | 1 |
| 12 | SN | 8 | 06.05.331 | S | 4 | 46 | NA | 0 | 0 | 1 |
| 12 | SN | 8 | 06.05.331 | N | 2 | 19.5 | 11 | 0 | 0 | 1 |
| 10 | SN | 7 | 06.07.322 | S | 1 | 25 | 75.5 | 0 | 0 | 1 |
| 13 | NS | 7 | 06.19.332 | N | 6 | 39 | 46 | 0 | 0 | 1 |
| 11 | NS | 6 | 06.06.324 | S | 4 | 46.5 | 52 | 0 | 0 | 1 |
| 2 | SN | 1 | 06.13.210 | N | 1 | 23.5 | NA | 0 | 0 | 1 |
| 2 | SN | 1 | 06.13.210 | N | 2 | 23.5 | NA | 0 | 0 | 1 |
| 2 | SN | 1 | 06.13.210 | N | 3 | 23.5 | NA | 0 | 0 | 1 |
| 2 | SN | 1 | 06.13.210 | N | 4 | 23.5 | NA | 0 | 0 | 1 |
| 4 | NS | 10 | 06.15.213 | N | 3 | 100 | 112 | 0 | 0 | 1 |
| 9 | NS | 5 | 06.05.321 | N | 3 | 7.5 | 44.5 | 0 | 0 | 1 |
| 9 | NS | 5 | 06.05.321 | N | 5 | 7.5 | 44.5 | 0 | 0 | 1 |
| 10 | SN | 7 | 06.07.322 | N | 1 | 44 | 43 | 1 | 1 | 1 |

| | | | | | | | | | | |
|---|---|---|---|---|---|---|---|---|---|---|
| 2 | SN | 1 | 06.13.210 | S | 1 | 18 | 57 | 1 | 1 | 1 |
| 13 | NS | 7 | 06.19.332 | N | 1 | 39 | 46 | 2 | 1 | 1 |
| 12 | SN | 8 | 06.05.331 | N | 4 | 19.5 | 11 | 0 | 0 | 2 |
| 12 | SN | 8 | 06.05.331 | N | 5 | 19.5 | 11 | 0 | 0 | 2 |
| 10 | SN | 7 | 06.07.322 | S | 6 | 25 | 75.5 | 0 | 0 | 2 |
| 10 | SN | 7 | 06.07.322 | S | 5 | 25 | 75.5 | 1 | 1 | 2 |
| 7 | SN | 8 | 06.14.232 | N | 2 | 23 | 57.5 | 4 | 1 | 2 |
| 4 | NS | 10 | 06.15.213 | S | 1 | 12.5 | NA | 0 | 0 | 3 |
| 7 | SN | 8 | 06.14.232 | N | 1 | 23 | 57.5 | 1 | 1 | 3 |
| 3 | NS | 8 | 06.13.211 | N | 3 | 0 | 39 | 0 | 0 | 1 |
| 3 | NS | 8 | 06.13.211 | N | 5 | 0 | 39 | 0 | 0 | 1 |
| 3 | NS | 8 | 06.13.211 | N | 6 | 0 | 39 | 0 | 0 | 1 |
| 1 | NS | 7 | 06.12.131 | N | 1 | 33 | 321 | 0 | 0 | 1 |
| 1 | NS | 7 | 06.12.131 | N | 2 | 33 | 321 | 0 | 0 | 1 |
| 3 | NS | 8 | 06.13.211 | S | 2 | 12 | 13 | 0 | 0 | 1 |
| 1 | NS | 7 | 06.12.131 | S | 2 | 63 | 3 | 0 | 0 | 1 |
| 10 | SN | 7 | 06.07.322 | S | 3 | 25 | 75.5 | 0 | 0 | 1 |
| 13 | NS | 7 | 06.19.332 | S | 6 | 16 | 18 | 0 | 0 | 1 |
| 5 | SN | 5 | 06.11.2213 | S | 1 | 42 | 173 | 0 | 0 | 1 |
| 5 | SN | 5 | 06.11.2213 | S | 5 | 42 | 173 | 0 | 0 | 1 |
| 11 | NS | 6 | 06.06.324 | N | 5 | 68 | 73 | 0 | 0 | 1 |
| 4 | NS | 10 | 06.15.213 | S | 6 | 12.5 | NA | 0 | 0 | 1 |
| 10 | SN | 7 | 06.19.322 | S | 4 | 32.5 | NA | 0 | 0 | 1 |
| 10 | SN | 7 | 06.19.322 | S | 1 | 32.5 | NA | 0 | 0 | 1 |
| 10 | SN | 7 | 06.19.322 | N | 6 | 5 | 9.5 | 0 | 0 | 1 |
| 3 | NS | 8 | 06.13.211 | N | 1 | 0 | 39 | 1 | 1 | 1 |
| 12 | SN | 8 | 06.05.331 | N | 1 | 19.5 | 11 | 1 | 1 | 1 |
| 5 | SN | 5 | 06.11.2213 | N | 2 | 54.5 | 52.5 | 1 | 1 | 1 |
| 11 | NS | 6 | 06.06.324 | S | 1 | 46.5 | 52 | 1 | 1 | 1 |
| 7 | SN | 8 | 06.14.232 | N | 6 | 23 | 57.5 | 1 | 1 | 1 |
| 6 | NS | 5 | 06.11.222 | S | 5 | 65.5 | 8 | 2 | 1 | 1 |
| 5 | SN | 5 | 06.11.2213 | S | 3 | 42 | 173 | 2 | 1 | 1 |
| 5 | SN | 5 | 06.11.2213 | S | 6 | 42 | 173 | 2 | 1 | 1 |
| 8 | NS | 7 | 06.06.311 | S | 5 | 7 | 65 | 2 | 1 | 1 |
| 10 | SN | 7 | 06.19.322 | N | 5 | 5 | 9.5 | 2 | 1 | 1 |
| 3 | NS | 8 | 06.13.211 | S | 3 | 12 | 13 | 3 | 1 | 1 |
| 6 | NS | 5 | 06.11.222 | S | 2 | 65.5 | 8 | 3 | 1 | 1 |
| 6 | NS | 5 | 06.11.222 | S | 4 | 65.5 | 8 | 3 | 1 | 1 |
| 11 | NS | 6 | 06.06.324 | S | 6 | 46.5 | 52 | 4 | 1 | 1 |
| 6 | NS | 5 | 06.11.222 | N | 1 | 44 | 4.5 | 9 | 1 | 1 |
| 1 | NS | 7 | 06.12.131 | S | 4 | 63 | 3 | 12 | 1 | 1 |
| 1 | NS | 7 | 06.12.131 | S | 5 | 63 | 3 | 19 | 1 | 1 |
| 12 | SN | 8 | 06.05.331 | N | 3 | 19.5 | 11 | 0 | 0 | 2 |
| 5 | SN | 5 | 06.11.2213 | S | 2 | 42 | 173 | 0 | 0 | 2 |
| 9 | NS | 5 | 06.05.321 | S | 2 | 39 | 305 | 0 | 0 | 2 |

| | | | | | | | | | | |
|---|---|---|---|---|---|---|---|---|---|---|
| 7 | SN | 8 | 06.14.232 | S | 4 | 5.5 | 44 | 0 | 0 | 2 |
| 4 | NS | 10 | 06.15.213 | N | 2 | 100 | 112 | 1 | 1 | 2 |
| 8 | NS | 7 | 06.06.311 | N | 3 | 142.5 | 81 | 1 | 1 | 2 |
| 10 | SN | 7 | 06.07.322 | S | 4 | 25 | 75.5 | 0 | 0 | 3 |
| 9 | NS | 5 | 06.05.321 | N | 4 | 7.5 | 44.5 | 0 | 0 | 3 |
| 4 | NS | 10 | 06.15.213 | N | 5 | 100 | 112 | 1 | 1 | 3 |
| 7 | SN | 8 | 06.14.232 | S | 1 | 5.5 | 44 | 7 | 1 | 3 |
| 9 | NS | 5 | 06.05.321 | N | 2 | 7.5 | 44.5 | 0 | 0 | 4 |
| 13 | NS | 7 | 06.07.332 | S | 6 | 4 | 24 | 0 | 0 | 4 |
| 12 | SN | 8 | 06.19.331 | S | 4 | 6.5 | 46.5 | 0 | 0 | 4 |
| 4 | NS | 10 | 06.15.213 | N | 6 | 100 | 112 | 1 | 1 | 4 |
| 13 | NS | 7 | 06.07.332 | N | 1 | 6 | 13.5 | 2 | 1 | 4 |
| 3 | NS | 8 | 06.13.211 | S | 5 | 12 | 13 | 0 | 0 | 2 |
| 3 | NS | 8 | 06.13.211 | N | 2 | 0 | 39 | 1 | 1 | 2 |
| 7 | SN | 8 | 06.14.232 | N | 5 | 23 | 57.5 | 1 | 1 | 2 |
| 3 | NS | 8 | 06.13.211 | S | 1 | 12 | 13 | 3 | 1 | 2 |
| 10 | SN | 7 | 06.07.322 | N | 2 | 44 | 43 | 3 | 1 | 2 |
| 7 | SN | 8 | 06.14.232 | N | 4 | 23 | 57.5 | 3 | 1 | 2 |
| 7 | SN | 8 | 06.14.232 | S | 6 | 5.5 | 44 | 5 | 1 | 2 |
| 6 | NS | 5 | 06.11.222 | N | 4 | 44 | 4.5 | 12 | 1 | 2 |
| 12 | SN | 8 | 06.05.331 | S | 3 | 46 | NA | 0 | 0 | 3 |
| 13 | NS | 7 | 06.07.332 | S | 3 | 4 | 24 | 0 | 0 | 3 |
| 10 | SN | 7 | 06.07.322 | N | 5 | 44 | 43 | 7 | 1 | 3 |
| 4 | NS | 10 | 06.15.213 | N | 4 | 100 | 112 | 0 | 0 | 4 |
| 13 | NS | 7 | 06.07.332 | S | 4 | 4 | 24 | 0 | 0 | 4 |
| 9 | NS | 5 | 06.05.321 | N | 1 | 7.5 | 44.5 | 1 | 1 | 4 |
| 7 | SN | 8 | 06.14.232 | S | 2 | 5.5 | 44 | 7 | 1 | 4 |
| 10 | SN | 7 | 06.19.322 | N | 1 | 5 | 9.5 | 3 | 1 | 5 |
| 13 | NS | 7 | 06.07.332 | N | 6 | 6 | 13.5 | 5 | 1 | 5 |
| 12 | SN | 8 | 06.19.331 | S | 1 | 6.5 | 46.5 | 1 | 1 | 6 |
| 12 | SN | 8 | 06.19.331 | S | 2 | 6.5 | 46.5 | 5 | 1 | 6 |
| 12 | SN | 8 | 06.05.331 | S | 6 | 46 | NA | 0 | 0 | 3 |
| 13 | NS | 7 | 06.19.332 | N | 3 | 39 | 46 | 0 | 0 | 3 |
| 13 | NS | 7 | 06.19.332 | N | 4 | 39 | 46 | 0 | 0 | 3 |
| 3 | NS | 8 | 06.13.211 | N | 4 | 0 | 39 | 1 | 1 | 3 |
| 3 | NS | 8 | 06.13.211 | S | 4 | 12 | 13 | 1 | 1 | 3 |
| 3 | NS | 8 | 06.13.211 | S | 6 | 12 | 13 | 1 | 1 | 3 |
| 10 | SN | 7 | 06.07.322 | S | 2 | 25 | 75.5 | 2 | 1 | 3 |
| 11 | NS | 6 | 06.06.324 | S | 3 | 46.5 | 52 | 2 | 1 | 3 |
| 1 | NS | 7 | 06.12.131 | N | 6 | 33 | 321 | 3 | 1 | 3 |
| 8 | NS | 7 | 06.06.311 | N | 4 | 142.5 | 81 | 7 | 1 | 3 |
| 13 | NS | 7 | 06.07.332 | N | 2 | 6 | 13.5 | 2 | 1 | 4 |
| 7 | SN | 8 | 06.14.232 | S | 3 | 5.5 | 44 | 2 | 1 | 4 |
| 7 | SN | 8 | 06.14.232 | S | 5 | 5.5 | 44 | 4 | 1 | 4 |
| 10 | SN | 7 | 06.07.322 | N | 4 | 44 | 43 | 1 | 1 | 5 |

| | | | | | | | | | | |
|---|---|---|---|---|---|---|---|---|---|---|
| 13 | NS | 7 | 06.07.332 | S | 5 | 4 | 24 | 0 | 0 | 6 |
| 12 | SN | 8 | 06.19.331 | S | 5 | 6.5 | 46.5 | 0 | 0 | 6 |
| 12 | SN | 8 | 06.19.331 | N | 4 | 31.5 | 31 | 5 | 1 | 6 |
| 12 | SN | 8 | 06.19.331 | S | 3 | 6.5 | 46.5 | 0 | 0 | 7 |
| 10 | SN | 7 | 06.07.322 | N | 3 | 44 | 43 | 0 | 0 | 4 |
| 9 | NS | 5 | 06.05.321 | S | 1 | 39 | 305 | 0 | 0 | 4 |
| 1 | NS | 7 | 06.12.131 | S | 3 | 63 | 3 | 2 | 1 | 4 |
| 10 | SN | 7 | 06.19.322 | N | 4 | 5 | 9.5 | 3 | 1 | 4 |
| 8 | NS | 7 | 06.06.311 | N | 5 | 142.5 | 81 | 3 | 1 | 5 |
| 10 | SN | 7 | 06.07.322 | N | 6 | 44 | 43 | 4 | 1 | 5 |
| 13 | NS | 7 | 06.07.332 | S | 2 | 4 | 24 | 0 | 0 | 6 |
| 7 | SN | 8 | 06.14.232 | N | 3 | 23 | 57.5 | 3 | 1 | 5 |
| 12 | SN | 8 | 06.19.331 | S | 6 | 6.5 | 46.5 | 7 | 1 | 6 |
| 12 | SN | 8 | 06.19.331 | N | 5 | 31.5 | 31 | 2 | 1 | 7 |
| 13 | NS | 7 | 06.19.332 | S | 3 | 16 | 18 | 11 | 1 | 9 |
| 13 | NS | 7 | 06.19.332 | S | 1 | 16 | 18 | 1 | 1 | 10 |
| 13 | NS | 7 | 06.19.332 | S | 2 | 16 | 18 | 10 | 1 | 10 |
| 13 | NS | 7 | 06.19.332 | S | 5 | 16 | 18 | 4 | 1 | 6 |
| 6 | NS | 5 | 06.11.222 | S | 3 | 65.5 | 8 | 5 | 1 | 6 |
| 13 | NS | 7 | 06.07.332 | N | 4 | 6 | 13.5 | 22 | 1 | 9 |
| 12 | SN | 8 | 06.19.331 | N | 3 | 31.5 | 31 | 21 | 1 | 10 |
| 12 | SN | 8 | 06.19.331 | N | 1 | 31.5 | 31 | 12 | 1 | 7 |
| 13 | NS | 7 | 06.07.332 | N | 5 | 6 | 13.5 | 4 | 1 | 9 |
| 12 | SN | 8 | 06.19.331 | N | 6 | 31.5 | 31 | 32 | 1 | 13 |
| 13 | NS | 7 | 06.19.332 | S | 4 | 16 | 18 | 5 | 1 | 8 |
| 13 | NS | 7 | 06.07.332 | S | 1 | 4 | 24 | 11 | 1 | 8 |
| 12 | SN | 8 | 06.19.331 | N | 2 | 31.5 | 31 | 15 | 1 | 15 |
| 13 | NS | 7 | 06.07.332 | N | 3 | 6 | 13.5 | 30 | 1 | 16 |
